# HomRank: Homogeneous RNA Ranking for 3D Structure Evaluation

**DOI:** 10.1101/2025.08.07.669087

**Authors:** Liang Li, Yinghua Yao, Ivona Martinović, Mandar Kulkarni, Yuangang Pan, Roland G. Huber, Mile Šikić

## Abstract

Leveraging rigorously curated RNA sequences and a novel ranking-based evaluation paradigm, we propose an improved pipeline for RNA 3D structure assessment. To enhance generalization, we construct a dataset comprising non-redundant single-chain RNA sequences from the PDB and apply unsupervised clustering to minimize data leakage. Based on this, we retrain the previous state-of-the-art method, ARES, to develop ARES+, which improves the top-1 success rate by 21% on the RNA Puzzles benchmark. To further boost near-native identification, we introduce HomRank, a homogeneous RNA ranking method that directly optimizes the selection of near-native conformations from candidate sets, resembling expert evaluation strategies. HomRank achieves 95% and 100% top-1 and top-5 success rates, respectively, significantly outperforming ARES+. These results demonstrate that carefully designed datasets and the expert-like selection paradigm can substantially improve the accuracy and robustness of RNA 3D structure assessment, offering a promising direction for deep learning-based RNA evaluation and near-native conformation selection.

---

RNA molecules play essential roles in diverse cellular processes, including gene expression, protein synthesis, and catalysis [1]. These functions are often governed by RNA’s intricate three-dimensional (3D) structures [2]. Although atomic-resolution RNA structures can be resolved through experimental techniques such as X-ray crystallography, nuclear magnetic resonance (NMR) spectroscopy, and cryogenic electron microscopy (Cryo-EM), these methods remain costly, labor-intensive, and low-throughput [3]. As a result, computational prediction of RNA 3D structures has emerged as a crucial alternative [4–7]. In contrast to the remarkable success of protein structure prediction—exemplified by AlphaFold [8] and RosettaFold [9]—RNA structure prediction still faces substantial challenges [10, 11]. These challenges stem from both limited training data—RNA structures account for less than 1% of entries in the Protein Data Bank [12]—and the intrinsic complexity of RNA folding, which involves dynamic, context-dependent energy landscapes [13]. To account for these uncertainties, most RNA structure predictors generate multiple candidate conformations per sequence, using different initializations or stochastic sampling strategies [14, 15]. While such ensembles help capture the conformational space, they introduce a fundamental challenge: *how to identify the most accurate RNA conformation among multiple candidates in the absence of the experimental ground truth*.

Traditional RNA 3D structure evaluation methods rely on knowledge-based statistical potentials, which benchmark candidates against the reference state that represents an idealized baseline conformation or energy state [16]. RASP [16] and 3dRNAscore [17] build all-atom distance potentials based on averaged reference states. *ɛ*-SCORE [18] focuses on relative arrangements and orientations of the nucleobases. DFIRE-RNA [19] introduces a distance-scaled statistical potential based on finite-ideal-gas reference state. Rosetta scoring function [20–22] incorporates a weighted combination of knowledge- and physics-based potentials, such as van der Waals interactions, base-pairing, and other statistical terms. rsRNASP [23] refines statistical potentials by distinguishing short- and long-range residue interactions. The recent cgRNASP [24] enhances rsRNASP by further refining the short-range interactions. Despite these advancements, a key limitation of this type of evaluation lies in the inherent difficulty of accurately modeling the reference state [25, 26].

Deep learning-based evaluation models bypass the need for manually defined reference states, enabling flexible, data-driven modeling. RNA3DCNN [27] uses 3D convolutional neural network to capture nucleotide interactions and local environments. lociPARSE [28] leverages the invariant point attention module from AlphaFold 2 [8] to extract geometric features of nucleotides. ARES [29] introduces E(3)-equivariant neural networks (E3NN) [30], which learn structural representations at the atomic level by enforcing equivariance, such that outputs transform consistently under symmetry operations applied to the input (e.g., rotations or translations), and invariance, whereby fundamental properties (e.g., energy) remain constant. By relying solely on atom types and three-dimensional coordinates—without incorporating RNA-specific priors such as base-pairing information—ARES has established itself as a widely used baseline.

Despite performing well on the RNA Puzzles benchmark [31–34], ARES essentially act as an RMSD regressor (left panel in Figure 1), using RMSD as a proxy for structure accuracy and enforcing a close match between the predicted score and the ground-truth RMSD. Such a manner has notable limitations: (1) Failing to capture relative structural differences: It focuses on scoring each candidate individually, omitting to discern relative structural difference among a batch of conformations for a specific sequence, i.e., homogeneous candidates. (2) Optimization imbalance due to RMSD variation: Candidates within a mini-batch arise from different sequences, i.e., heterogeneous RNAs, with varying RMSD ranges. Longer-chain RNAs typically exhibit broader ranges [35], which can lead to training errors dominating the optimization. (3) Sensitivity to outliers: Regression task is inherently vulnerable to outliers, degrading the robustness. These limitations hinder ARES’s ability to generalize effectively to new data, particularly when it differs significantly from the training data.

**Fig. 1:**
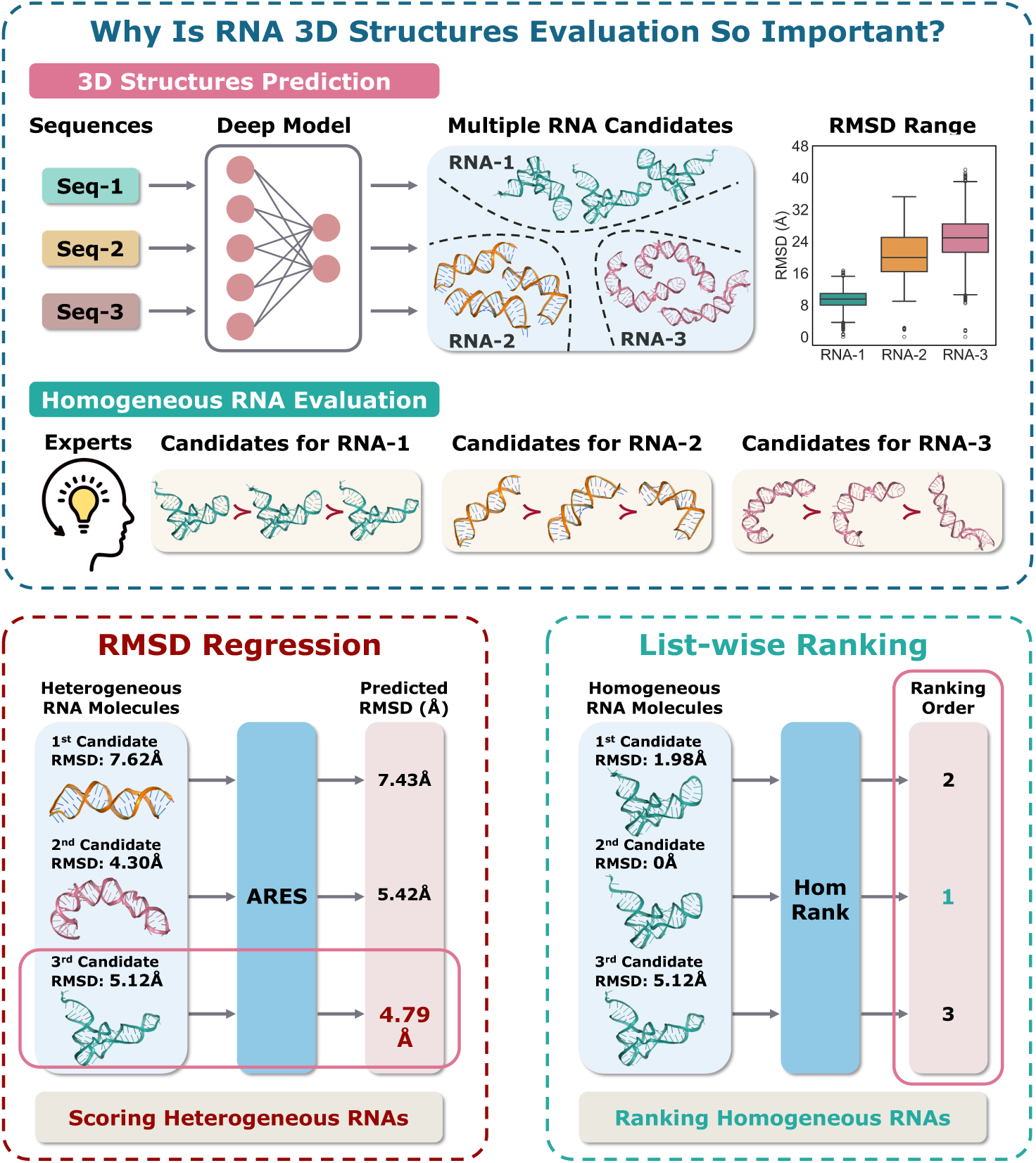
RNA 3D Structure Evaluation. (Top) Motivation for developing robust evaluation models; (bottom left) conventional RMSD-based regression approach; (bottom right) our proposed *HomRank* paradigm. Traditional methods, such as *ARES*, train RMSD regressors to score each candidate independently, failing to capture relative structural differences among conformations. In contrast, *HomRank* employs a listwise ranking framework that explicitly models these differences across structurally similar candidates, closely aligning with expert-driven evaluation practices.

Therefore, instead of following the conventional RMSD regression, we propose a novel homogeneous RNA ranking technique, termed HomRank, as shown in the right panel of Figure 1, inspired by the observation that experts rank homogeneous structures based on domain expertise (top panel of Figure 1). Specifically, we introduce a list-wise ranking technique [36] to directly model the relative structural difference of homogeneous candidates, rather than predicting absolute RMSD scores individually, which effectively addresses the above issues of RMSD regression. On the other hand, to ensure model generalization, we construct a new diverse dataset with rigorous splitting. We utilize the Bio.Align module from the BioPython library [37] to align RNA sequences and calculate their similarity, followed by hierarchical clustering to group them in an unsupervised manner. The dataset is split according to automatically identified clusters, rather than relying on manual splitting [27] or the publication years of RNAs [28, 29], thereby effectively eliminating human interference and minimizing the risk of potential data leakage. Extensive experiments demonstrate that HomRank achieves notable improvements in scoring accuracy compared to RMSD regression models, and effectively captures fine-grained biological characteristics without explicit specification in the model design. Furthermore, HomRank exhibits promising potential in distinguishing different RNA conformations from various deep learning-based RNA 3D structure prediction models, highlighting its effectiveness as a general-purpose RNA 3D structure evaluation model, regardless of whether the structures originate from a specific or different RNA 3D structure prediction models.

## Results

HomRank adopts the E3NN architecture and equips with the ListNet ranking technique [36], as detailed in Methods. To enable rigorous evaluation of RNA 3D structure scoring models, we construct a diverse dataset by collecting non-redundant single-chain RNAs from the PDB database and applying pairwise sequence alignment followed by hierarchical clustering. This process yields three distinct clusters (Fig. 2): CLS-0, CLS-1, and CLS-2, where CLS-0, the largest, is used for training^1^, and CLS-1 and CLS-2 serve as TestCLS-1 and TestCLS-2 to test generalization. We also test our performance on the RNA Puzzles benchmark [34]. We retrain the ARES model on the new training dataset, termed as ARES+. To check whether HomRank can refine the pre-trained RMSD regression model, we design a control group experiment: HomRank-PT, which is identical in design with HomRank while incorporating a pre-training phase with a MSE-based RMSD regression loss as ARES.

**Fig. 2:**
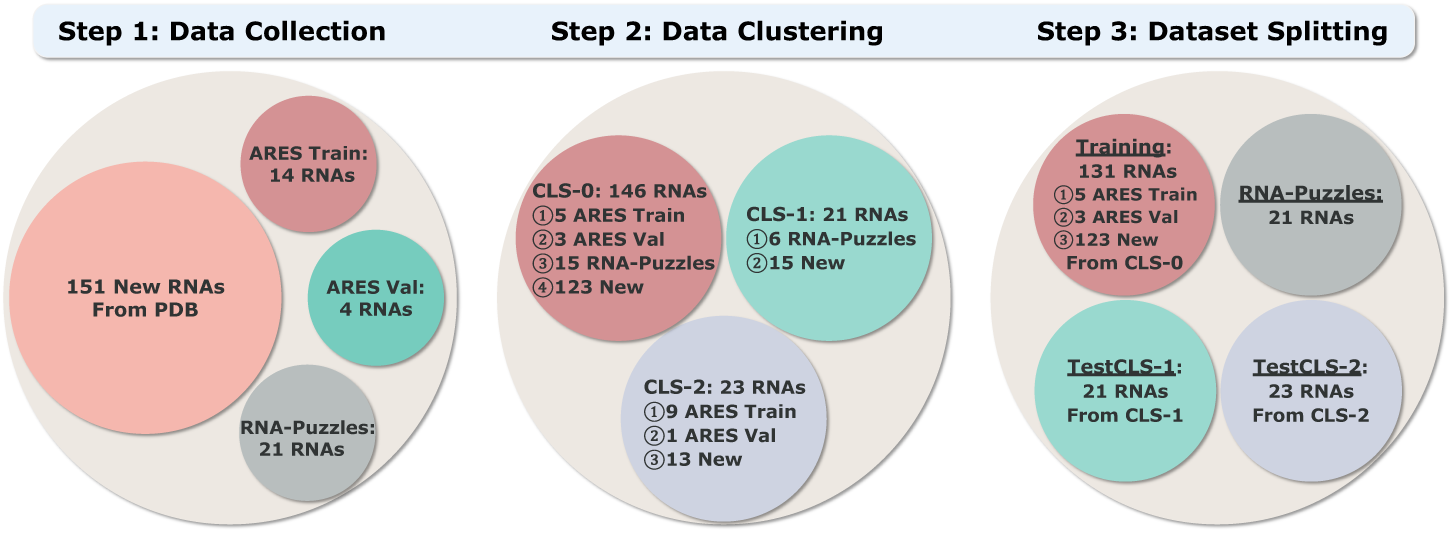
Workflow of meticulous data collection, unsupervised clustering, and rigorous splitting. The three clusters, CLS-0, CLS-1, and CLS-2, are automatically identified through hierarchical clustering.

### Enhanced Performance with HomRank Powered by a New Dataset

#### Evaluating RNA Puzzles Benchmark

Figure 3(a)-(c) illustrates the overall comparison on the popular RNA Puzzles benchmark, which consists of 21 RNAs from the RNA Puzzles structure prediction challenge [34]. As noted in [29], the structural candidates were generated by Rosetta FARFAR2 sampling method [22]. Each RNA in this benchmark comprises at least 1,500 structural candidates, including one native conformation, 1% near-native structures (RMSD ≤ 2Å), and the remaining as decoy candidates (RMSD *>* 2Å).

**Fig. 3:**
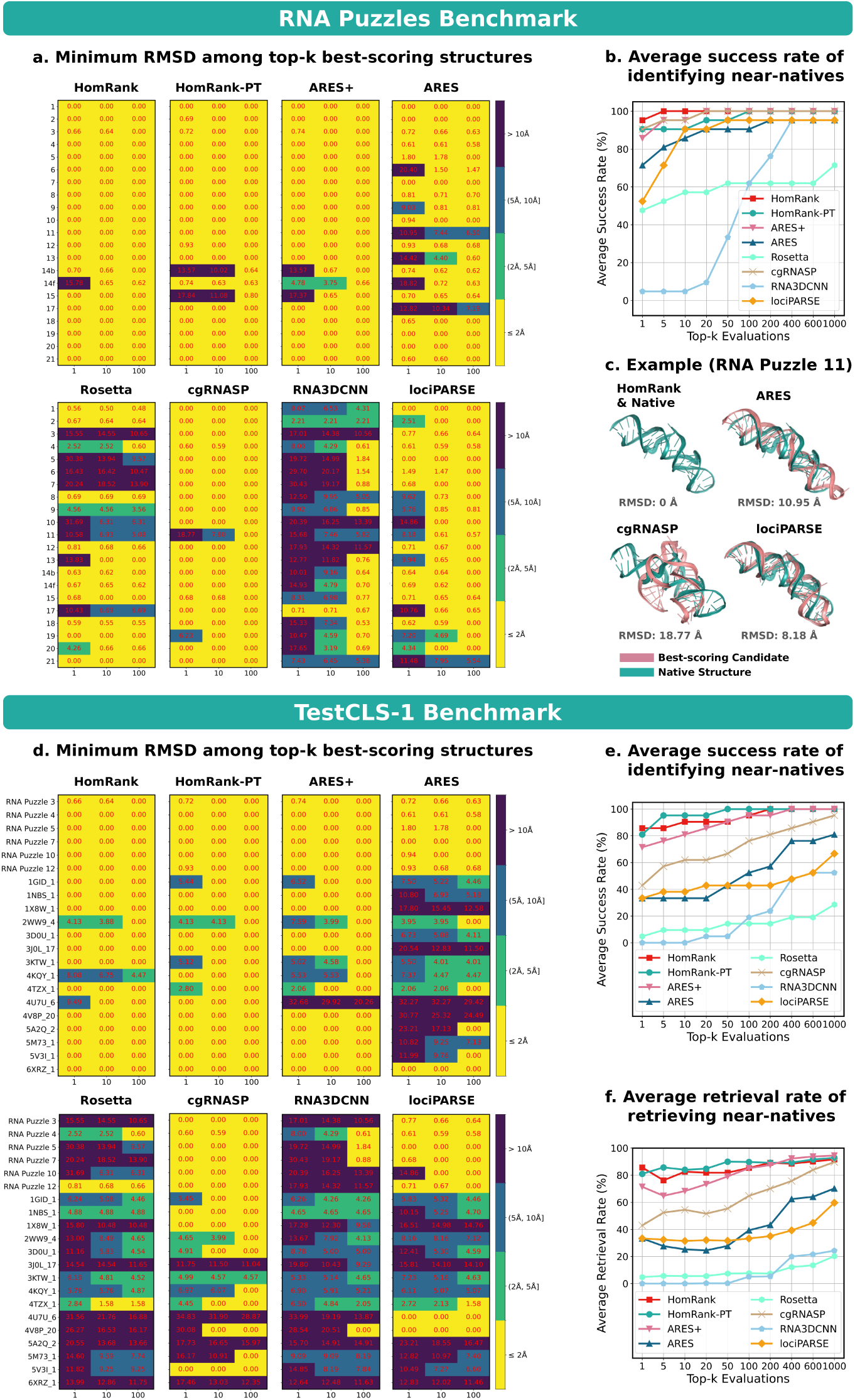
HomRank achieves superior performance in identifying near-native candidates (RMSD ≤ 2Å) on RNA Puzzles benchmark (a-c) and Test CLS-1 benchmark (d-f). **a.** Minimum RMSD of the top 1, 10, 100 best-scoring RNA candidates. RMSD values (highlighted in red) are categorized as follows: ≤ 2Å (yellow), (2Å, 5Å] (green), (5Å, 10Å] (blue), and above 10Å (purple). **b.** Average success rate of identifying near-native structures across all RNAs for the top-*k* evaluations. **c.** Case study on RNA Puzzle 11. **d.** Minimum RMSD for the top 1, 10, 100 best-scoring candidates. **e.** Average success rate of identifying near-native structures across all RNAs for the top-*k* evaluations. **f.** Average retrieval rate of retrieving near-native structures across all RNAs for the top-*k* evaluations. Metric definitions are provided in Methods.

Figure 3(a) depicts the minimum RMSD among the top 1, 10, 100 best-scoring candidates. HomRank achieves the strongest performance in identifying near-native candidates, followed by HomRank-PT and ARES+. The improvements of ARES+ over ARES indicate the benefits conferred by new training dataset. Among the five baseline methods, cgRNASP emerges as the most competitive, whereas RNA3DCNN exhibits the worst performance.

Figure 3(b) presents the average success rate of identifying near-natives in the top-*k* evaluations. HomRank identifies 95% near-natives in the top-1 evaluations, outperforming HomRank-PT and cgRNASP (both at 90%), as well as ARES+ (86%), ARES (71%), lociPARSE (52%), Rosetta (48%), and RNA3DCNN (5%). Furthermore, HomRank is the only method to achieve a 100% average success rate within the top-5 evaluations, surpassing ARES+ and cgRNASP, which require top-20 evaluations, as well as HomRank-PT, which needs top-100 evaluations to reach a comparable success rate. In contrast, ARES, Rosetta, RNA3DCNN, lociPARSE fail to achieve 100% even when the search range is expanded to the top-1000 candidates, with their metrics remaining between 71% and 95%, indicating their limited scoring capacity.

In the case study of RNA Puzzle 11 shown in Figure 3(c), HomRank is compared with ARES and two leading competitors, cgRNASP and lociPARSE. HomRank accurately identifies the native conformation as the best-scoring candidate, while the baselines erroneously select decoy candidates with significantly higher RMSD values: ARES (RMSD=10.95Å), cgRNASP (RMSD=18.77Å), and lociPARSE (RMSD=8.18Å). These results highlight the superior scoring accuracy of HomRank on the RNA Puzzles benchmark. Additional experimental results on the method comparisons are provided in Supplementary Fig. S2.

#### Evaluating TestCLS-1 Benchmark

Model generalization is crucial for the RNA 3D structure evaluation task. Although existing deep learning-based scoring methods perform well on in-distribution data, they may exhibit degraded accuracy on new RNAs, hidering practical usage of such tools [38]. Figure 3(d)-(f) compares the performance on the TestCLS-1 benchmark, which comprises 21 RNAs belonging to CLS-1 – a cluster distinct from the training data, providing a rigorous evaluation of model generalization on unseen structures. Figure 3(a) presents the minimum RMSD for the top 1, 10, 100 best-scoring candidates. HomRank, HomRank-PT, and ARES+ consistently outperform the compared baselines. ARES+ demonstrates significant improvements over ARES, highlighting the contributions of our dataset in enhancing model generalization. HomRank-PT achieves further improvements, validating our technical contribution that homogeneous RNA ranking effectively enhances top-*k* retrieval performance. For the compared baselines, cgRNASP is the strongest competitor. However, its performance on TestCLS-1 benchmark is significantly inferior to that on RNA Puzzles benchmark, indicating its limited generalization and adaptability on unseen RNAs. Figure 3(b) quantifies the average success rate of identifying near-natives within the top-*k* evaluations, assessing the capacity to identify near-native conformations. HomRank, HomRank-PT, and ARES+ identify 86%, 81%, and 71% of near-native molecules in the top-1 evaluations, significantly surpassing cgRNASP (43%), ARES (33%), lociPARSE (33%), Rosetta (5%), and RNA3DCNN (0%). Besides, HomRank, HomRank-PT, and ARES+ achieve 100% average success rate within the top-100, top-50, and top-100 evaluations, respectively. However, even when we extend the search range to the top-1000 candidates, the metric for the five baselines ranges from 29% to 95%.

Figure 3(c) depicts the average retrieval rate of near-natives, measuring their proportion within the top-*k* evaluations. We observe that HomRank, HomRank-PT, and ARES+ consistently exceed the baselines by substantial margins, demonstrating superior retrieval accuracy and reliability.

In summary, the two knowledge-based baselines and three recent deep learning baselines struggle to generalize on the challenging TestCLS-1 benchmark. ARES+ demonstrates notable improvement over ARES, highlighting the enhanced performance powered by the new data. Furthermore, HomRank and HomRank-PT show promising capacity in retrieving near-native candidates, validating the effectiveness of homogeneous RNA ranking technique. More experimental results on TestCLS-1 are available in Supplementary Figure S3, while those on TestCLS-2 are can be found in Supplementary Figure S4.

### Ranking Pairs with Varying Structural Similarities

This part investigates model performance in ranking pairs with varying structural similarity measured by ΔRMSD, inspired by a practical scenario faced by structural biologists: when comparing pairs with large ΔRMSD, distinguishing the candidate closer to the native conformation is relatively straightforward. However, as ΔRMSD decreases, structural similarities increase, making the judgment more challenging and raising the likelihood of mistakes.

We select twelve RNAs from the TestCLS-1 and TestCLS-2 and define six initial ΔRMSD intervals: (0Å, 2Å], (2Å, 4Å], (4Å, 6Å], (6Å, 8Å], (8Å, 10Å], and ≥ 10Å. These intervals are adaptively adjusted based on the ground-truth RMSD values of RNA candidates. For each interval, we randomly sample 1,000 pairs and calculate accuracy, i.e., the proportion of correct pairwise rankings.

Figure 4 illustrates that for most models, the ranking accuracy improves as ΔRMSD increases, similar to the evaluation paradigm of domain experts. HomRank, HomRank-PT, and ARES+ consistently achieve higher accuracy than other baselines in most cases. An interesting finding is that Rosetta and lociPARSE show a negative correlation in the case of 2WW9 4, with accuracy dropping from approximately 50% for ΔRMSD ∈ (0Å, 2Å] to around 20% for RMSD ≥ 10Å. Similarly, for 4U7U 6, the accuracy of cgRNASP and ARES declines sharply, from 40% for ΔRMSD ∈ (0Å, 2Å] to below 10% for ΔRMSD ≥ 10Å. These results suggest that Rosetta, lociPARSE, cgRNASP, and ARES have limited capacity to generalize to cases with significant structural discrepancies, likely due to biases in the training data or inherent deficiency in capturing critical structural features.

**Fig. 4:**
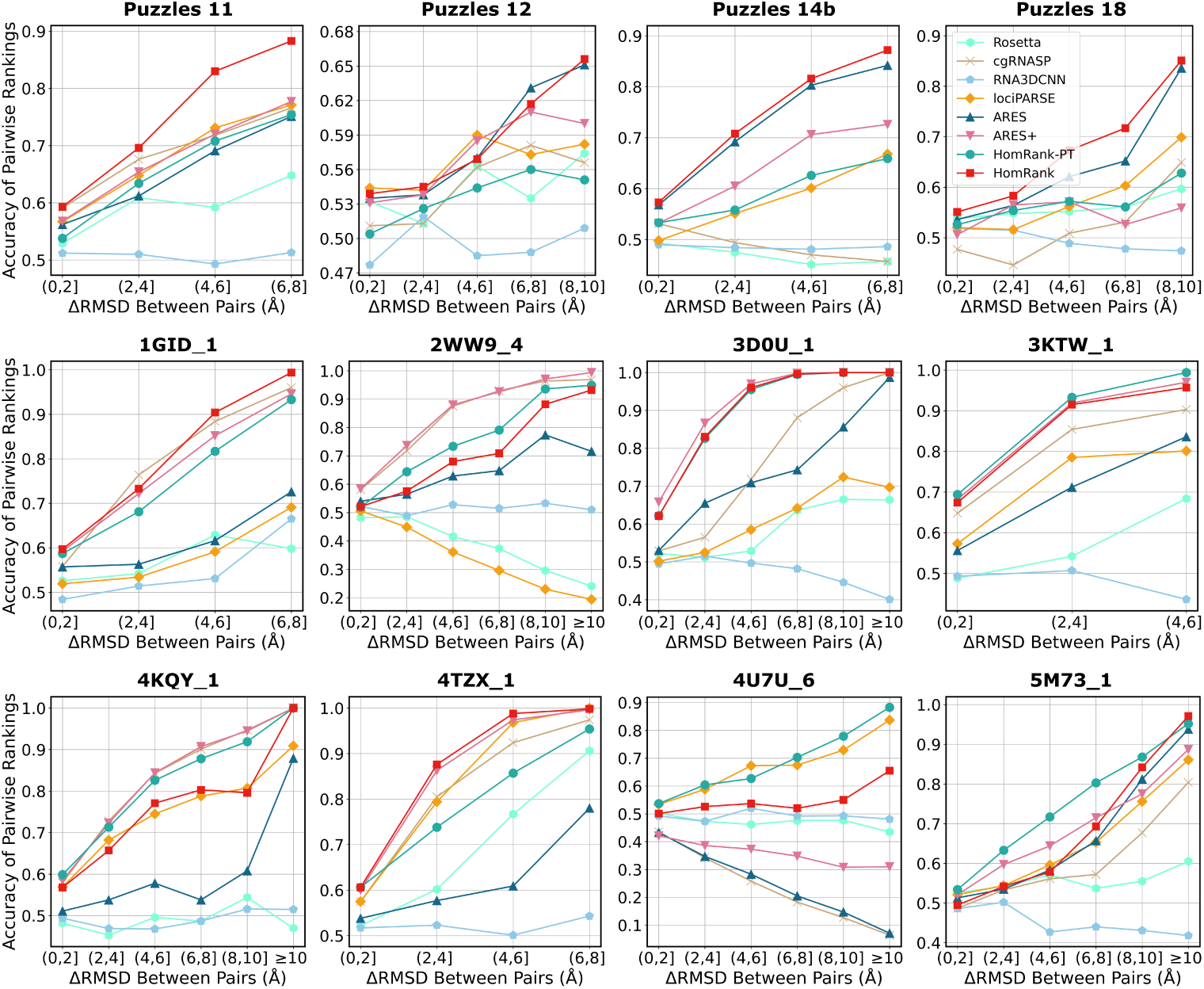
Ranking accuracy versus ΔRMSD.

Another observation is that incorporating homogeneous RNA ranking into the training process effectively mitigates negative correlation and improves ranking accuracy. An evidence is the case of 4U7U 6, where ARES exhibits a steep drop in accuracy as ΔRMSD increases, ARES+ improves the performance over ARES but still retains a negative correlation. In contrast, HomRank-PT not only surpasses ARES and ARES+ with higher accuracy but also reverses the trend, achieving a positive correlation. These results demonstrate the effectiveness of homogeneous RNA ranking in capturing key structural features that differentiate high-fidelity candidates from the low-fidelity ones, enabling models to reliably identify the candidate closer to the native conformation.

### Identifying Key Characteristics Within RNA Double Helices

We follow [29] to introduce RNA double helices with varying inter-strand distances, shown in Figure 5(a), to investigate whether the models can identify key biological characteristics without explicit specification in the model.

**Fig. 5:**
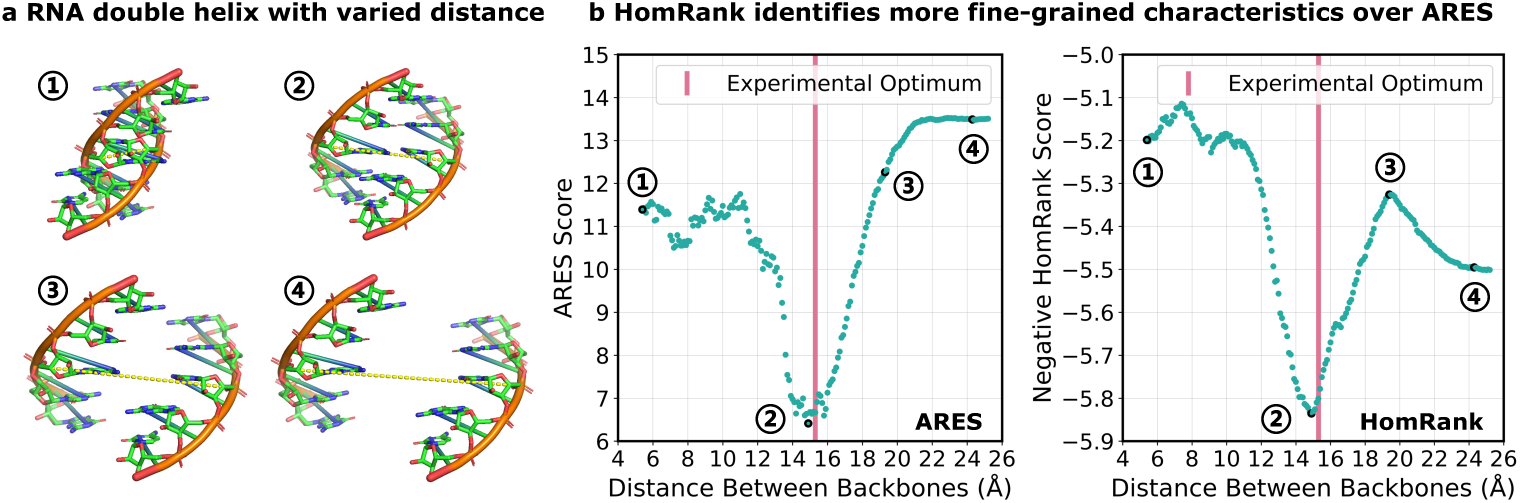
HomRank captures finer-grained structural characteristics without prior specification. **a**. A case of RNA double helices, where the inter-strand distance is measured between the C-4’ atoms of the central base pair (yellow dotted lines). **b**. With varying inter-strand distance, both HomRank and ARES assign the best scores when the distance closely aligns with the experimentally determined optimal distance (pink vertical line). However, ARES fails to capture the finer-grained transitions from the increasing dominance of attractive forces to diminishing interactions (point 2 to 4). For clarity, we take the negative ranking score for HomRank, as higher ranking scores correspond to higher-fidelity conformations.

Figure 5(b) compares the results of HomRank and ARES. For convenience, we plot the negative ranking score for HomRank. Both models accurately assign lowest score when the inter-strand distance of the RNA double helices approaches the experimentally determined optimal distance (red vertical line), corresponding to ideal base pairing. As the inter-strand distance decreases below the equilibrium distance (the seconde point), both HomRank and ARES assign higher scores. This behavior is attributed to significantly increased repulsive interactions at short-range distance. Conversely, when the distance exceeds the optimal one, HomRank exhibits distinct behavior depending on the distance of separation. Specifically, its negative score firstly increases, reaching a peak (the third point), then decreases and eventually stabilizes at a distant separation (the fourth point). The rising phase can be explained by the growing dominance of attractive forces, whereas the subsequent falling phase occurs as the strands move too far apart for significant interactions, resulting in a rapid decline of attractive forces. In the stable phase, the distance is sufficiently large to render interactions negligible. However, these finer-grained characteristics, particularly the transitions within the curve, are not captured by ARES. These findings indicate HomoRank’s superior sensitivity to the intricate interplay of structural and interaction dynamics in RNA double helices across varying inter-strand distance.

### Scoring Predictions from Six Deep Learning-based RNA 3D Structure Prediction Models

This part evaluates whether HomRank can generalize to discriminate the predictions of different deep learning-based RNA 3D structure prediction models, a challenging task due to structural variability and prediction uncertainties across different prediction methods, and the significant discrepancy from the training set in this work.

We introduce a testing benchmark (Dataset 3) from the recent review [39] on deep learning-based RNA 3D structure prediction models, which consists of 84 RNA sequences, each with the predictions from six state-of-the-art RNA 3D structure prediction models: DRfold [14], DeepFoldRNA [40], RhoFold [41], RoseTTAFoldNA [42], trRosettaRNA [6], and AlphaFold 3 [43]. Figure 6(a) shows the boxplot of the RMSD distribution of their predictions for the 84 sequences. We observe that RMSD values are centered around a median of 10Å, ranging from 9.64Å (DeepFoldRNA) to 16.33Å (AlphaFold 3). Especially, all six prediction models struggled with several RNA sequences, where the RMSD values of their predictions reached 75Å, far exceeding the pursuit of structural biologists that achieve near-native predictions (RMSD ≤ 2Å). First, we assess the retrieval performance in identifying the best predictions with the lowest RMSD. Figure 6(b) presents the boxplot of the RMSD distribution of the best-scoring structures, chosen from six predictions for each of the 84 RNA sequences. The “Optimal” category denotes a hypothetical perfect scoring model that precisely select the best predictions, defining the upper bound that scoring models strive to approximate. We observe that: (1) HomRank demonstrates the closest performance to the optimal baseline, as evidenced by its significantly narrower RMSD distribution and lower median RMSD, outperforming other scoring models. (2) ARES and ARES+ produce nearly the same results, indicating that retraining ARES on the new dataset does not improve its ability to distinguish predictions from different prediction models. (3) HomRank-PT achieves a lower median RMSD than ARES+ but exhibits a broader RMSD range, suggesting that its capacity for distinguishing structures generated by different models remains limited. (4) Among the compared models, Rosetta demonstrates the most competitive performance, followed by lociPARSE and RNA3DCNN, while cgRNASP exhibits the weakest result.

**Fig. 6:**
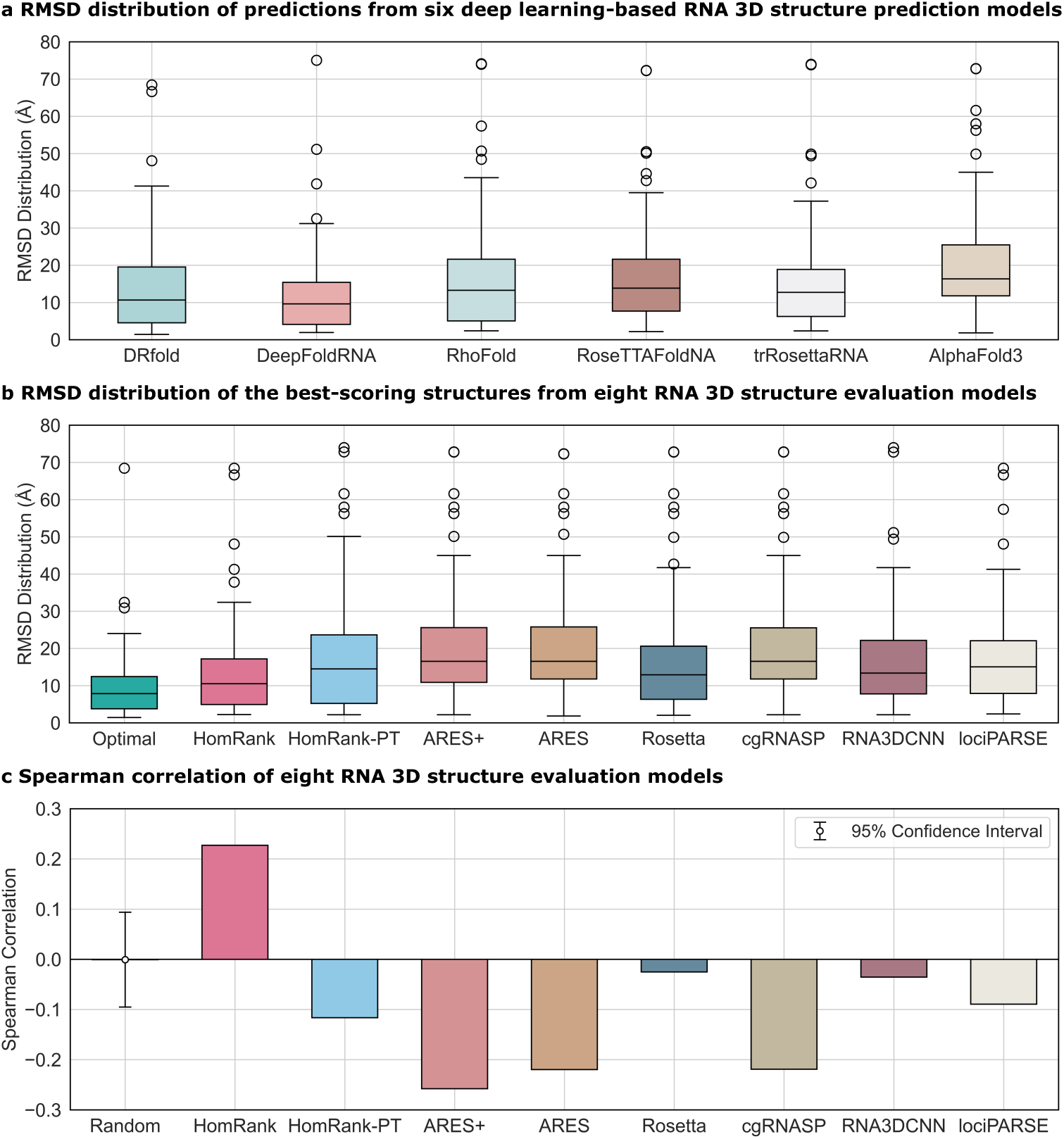
HomRank demonstrates promising capacity in ranking structures from six deep learning-based RNA 3D structure prediction models. **a**. Boxplot showing the RMSD distribution of predictions from six deep learning-based RNA 3D structure prediction models. **b**. Boxplot illustrating the RMSD distribution of the best-scoring structures from eight RNA 3D structure evaluation models. The “Optimal” category denotes a theoretical perfect scoring function that precisely identifies the best structure with the minimal RMSD among the six predictions. **c**. Spearman correlation of eight RNA 3D structure evaluation models. The “Random” category represents the result of random ranking and serves as a reference baseline.

Furthermore, we compute the Spearman correlation coefficients between the RMSD values and the predicted scores to assess how well the scoring model ranks the predictions in agreement with the true ranking, as shown in Figure 6(c). As a reference, we also include the result of random ranking, computed over 10,000 independent trials, which generates a mean Spearman correlation coefficient of -0.0008, with a 95% confidence interval ranging from -0.0952 to 0.0939. We observe that HomRank is the only one to achieve a positive Spearman correlation (0.2272), whereas all other scoring models exhibit correlations ranging from -0.2578 to -0.0252, performing even worse than random ranking. The unsatisfactory performance of most scoring models can be attributed to the change of the testing setting from that of the training process. In particular, most methods (except lociPARSE) are trained on data from a single prediction method, while the testing task requires them to score candidates from different prediction models – a significantly more complex and challenging task. Although lociPARSE is trained on a dataset consisting of structures from five deep learning-based and two non-deep RNA 3D prediction models, its performance remains limited. In contrast, HomRank demonstrates promising performance in distinguishing RNA conformations generated from various RNA 3D structure prediction models. This improvement is mainly due to our ranking-based optimization design. By focusing on distinguishing structural differences among RNA candidates, HomRank exhibits greater robustness and generalization on scoring conformations from different RNA 3D structure prediction models. Although HomRank’s performance still leaves room for improvement, it demonstrates promising potential as a general-purpose RNA 3D structure scoring model for assessing RNAs, regardless of whether they are from a single RNA 3D structure prediction model or multiple prediction models.

## Discussion

This paper provides a comprehensive improvement over previous works in RNA 3D structure evaluation, including providing a new RNA dataset for rigorous evaluation of model generalization, and proposing a novel scoring paradigm for enhancing the accuracy. The new dataset consists of diverse, non-redundant, single-chain RNAs from PDB, coupled with unsupervised data grouping and splitting, facilitating a rigorous evaluation of model generalization. As a representative RMSD regression-based scoring method, the ARES model retrained on the new training set, i.e., ARES+, demonstrates significantly improved top-*k* retrieval accuracy compared to its original version. Furthermore, we propose HomRank, a novel scoring paradigm designed to directly learn the relative structural differences among homogeneous RNA candidates. Unlike traditional RMSD regression-based scoring methods, HomRank emphasizes list-wise rankings, better aligning with the evaluation paradigm adopted by biological experts. We also develop HomRank-PT, a variant of HomRank that is identical in design but incorporates a pre-training phase using an MSE-based RMSD regression loss. Both HomRank and HomRank-PT outperform state-of-the-art knowledge-based methods and deep learning-based baselines, exhibiting promising accuracy and generalization. Furthermore, HomRank demonstrates superiority in distinguishing RNA conformations from various deep learning-based RNA 3D structure prediction models, which is a challenging task due to structural variability and prediction uncertainties from different prediction methods. While there is still room for improvement, HomRank shows strong potential as a general-purpose scoring model for assessing the quality of RNA 3D structures, no matter whether they are from a specific or different prediction models.

Despite the success demonstrated here, the overall performance of RNA scoring models remains limited, particularly evident on the TestCLS-2 benchmark, where all the baselines, including ours, show room for improvement in scoring accuracy. This limitation likely stems from the scarcity of experimentally resolved RNA structures, which restricts the models to generalize to unseen data. For our HomRank, its limitation is reliance on the ARES backbone. While effective, incorporating more high-capacity architectures, such as transformer-based architecture, has the potential to further enhance both the generalization and accuracy, which will be our future work. Nevertheless, as a proof of concept, HomRank provides a flexible and useful platform to the community for RNA 3D structure evaluation.

## Methods

### Data Collection, Clustering, and Splitting

Figure 2 shows the data processing pipeline, including data collection, clustering, and splitting.

In the data collection stage, we exclusively collect a comprehensive set of 190 non-redundant and single-chain RNA molecules, including 151 non-redundant RNAs from the PDB database, along with RNA Puzzles data (21 RNAs) and those from ARES: training (14 RNAs) and validation (4 RNAs).

During the data clustering stage, we apply the Bio.Align module [37] to perform pairwise sequence alignment for 190 RNAs and calculate their similarity. Using the resulting similarity map, we apply hierarchical clustering, which categorize them into three distinct groups: a larger cluster (CLS-0) comprising 146 RNAs, and two smaller clusters (CLS-1 contains 21 RNAs, CLS-2 includes 23 RNAs). A detailed clustering map is shown in Supplementary Figure S1.

The dataset splitting is based on the clustering results. CLS-0 is designated as the training set due to its larger size than the other clusters, while CLS-1 and CLS-2 are assigned as benchmarks, referred to as TestCLS-1 (21 RNAs) and TestCLS-2 (23 RNAs), respectively. It is worth noting that the 15 RNA Puzzles within CLS-0 are excluded from the training data, as these are commonly recognized as standard benchmarks [22, 29]. Consequently, the training set comprises 131 RNAs. Consistent with prior works [28, 29], the RNA Puzzles (21 RNAs) are set as an independent benchmark. Table 1 gives the details of data splitting.

**Table 1:** Details of dataset splitting.

| Dataset | Training | TestCLS-1 | TestCLS-2 | RNA Puzzles |
| --- | --- | --- | --- | --- |
| RNA Molecules | 131 | 21 | 23 | 21 |
| Sequence Length Range | 22-144 | 61-257 | 12-49 | 41-188 |
| Structural Candidates | 131,000 | 152,917 | 23,000 | 451,629 |

An interesting observation is that 5 training and 3 validation RNAs from ARES, along with 15 RNA Puzzles, are clustered into CLS-0. Similarly, CLS-1 contains 9 training and 1 validation RNAs from ARES. These findings indicate a critical limitation of conventional splitting methods, specifically manual partitioning [27] or reliance on publication years of RNA data [28, 29], may unintentionally introduce data leakage between training and testing data, thereby impairing the reliability of model evaluation.

For the 151 new RNAs, each molecule includes 1,000 candidates generated using molecular dynamics simulations. The structural candidates from ARES (1,000 for each of 18 RNAs) and RNA Puzzles benchmark (total 451,630 for 21 RNAs) from [29] remain unchanged. To evaluate the retrieval capacity for near-native structures (RMSD ≤ 2Å w.r.t. the experimentally resolved native structure), approximately 1% of the candidates for each RNA in the benchmarks are near-natives, while the rest are decoy structures (RMSD *>* 2Å). Details are provided in Supplementary Table S1, Table S2 and Table S3.

### RMSD Regression vs. Homogeneous RNA Ranking

This section compares the conventional RMSD regression and HomRank.

#### RMSD Regression

Conventional deep learning-based scoring models typically use RMSD as a proxy for structural accuracy, training a regressor to predict scores that closely approximate the ground-truth RMSD [29, 44]. RMSD quantifies the structural deviation of candidates from the native conformation, with lower RMSD indicating closer alignment. Let 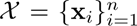 be the set of training data, and let 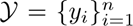 denote the corresponding ground-truth RMSD values. The training objective is to minimize the Mean Squared Error (MSE) loss, defined as:

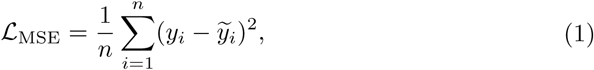

where *y_i_* is the ground-truth RMSD for the *i*-th instance, and 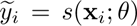 is the predicted RMSD. Here, *s*(·; *θ*) denotes the regression score function modeled by a neural network parameterized by *θ*, which is optimized during training.

#### Homogeneous RNA Ranking

First, let us consider how domain experts evaluate RNA 3D structures. Given a set of candidates for a specific sequence, i.e., homogeneous candidates, denoted as 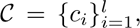 biological experts leverage their domain knowledge (e.g., base pairing patterns, tertiary interactions, and overall stability) to identify candidates with favorable characteristics, such as energetically stable conformations. Let *s*(·) represent a scoring function that encapsulates experts’ assessments, which reflects their preferences for these candidates. Such an expert-driven evaluation can be formulated as:

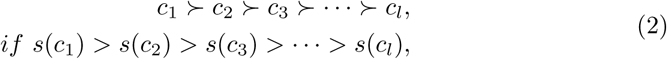

where *c*_1_ ≻ *c*_2_ denotes the candidate *c*_1_ is superior to *c*_2_, and *l* is the length of mini-list, which can be adjusted flexibly based on specific requirements or preference. To align with expert-driven evaluation, we adopt ListNet [36], a list-wise ranking technique, to optimize the rankings of homogeneous candidates. In this setting, *s*(**x***_i_*; *θ*) represents the ranking score function modeled by a neural network parameterized by *θ*, where higher scores indicate a closer alignment with the native conformation. Given *l* homogeneous candidates, each permutation *π* ∈ 𝒢*_l_* (representing a mini-list consisting of *l* candidates) is associated with a probability as follows,

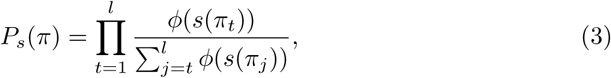

where 𝒢*_l_* is the set of all permutations of *l* candidates, *π_t_* is the *t*-th element of *π*, and *ϕ*(·) is a strictly positive, monotonically increasing transformation, which often uses the softmax function 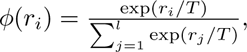 where *r* = [*r*_1_, *r*_2_, · · ·, *r_l_*] is a list of scalar scores and *T* is the temperature coefficient. The softmax function maps ranking scores to a probability distribution, naturally modeling the likelihood of different permutations. Additionally, exponential transformation amplifies score differences, enabling higher scores to have a stronger influence on the ranking probabilities.

It is evident that *P_s_*(*π*) ≥ 0 and 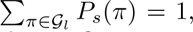 confirming that *P_s_*(*π*) defines a valid probability distribution over the set 𝒢*_l_*. However, since there are *l*! permutations, the computation becomes intractable for large *l*. To improve efficiency, it is common to consider the top-*k* probability, where the top-*k* candidates in the permutation *π* are specified (denoted as 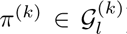). This method reduces the computation from 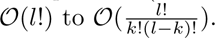 Eq. 3 is transfo*^l^* rmed into

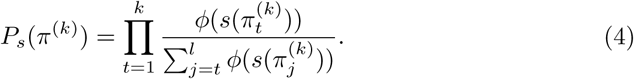

Also, *P_s_*(*π*^(^*^k^*^)^) ≥ 0 and 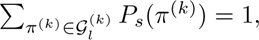 ensuring that the top-*k* probabilities form a valid probability distribution over the set 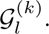 In practice, we often set *k* = 1 to prioritize the top-1 candidate, as the goal of RNA 3D structure evaluation is to maximize the likelihood of the most accurate candidate being ranked first. This manner further reduces the computation to 𝒪(*l*).

The above describes the construction of the predicted ranking distribution. For the ground-truth ranking distribution, a similar technique is applied. The difference is that the ground-truth RMSD values are not directly used but are instead converted into ranking scores (e.g., 1, 1/2, · · ·, 1/l), through a reciprocal transformation of the ranking order determined by the ground-truth RMSD, where smaller RMSD values correspond to higher rankings. Additionally, candidates with ΔRMSD ≤ *δ*Å are assigned the same ranking order to account for the negligible structural deviations, where *δ* is a tunable hyper-parameter.

Naturally, HomRank loss is defined as the cross-entropy loss, measuring the alignment between the ground-truth and predicted ranking distributions, which is expressed as:

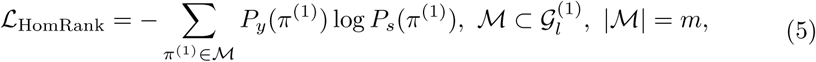

where *P_y_*(*π*^(1)^) is the top-1 ground-truth ranking distribution, *P_s_*(*π*^(1)^) is the corresponding predicted ranking distribution, ℳ is a subset of 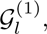 consisting of *m* mini-lists *π*^(1)^ randomly sampled from 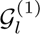 within each mini-batch.

The advantages of HomRank are as follows: (1) Alignment with expert-driven evaluation paradigm: HomRank focuses on ranking homogeneous candidates from the same sequence, aligning closely with expert-driven evaluation and the inference task in RNA 3D structure evaluation. (2) Improved scoring accuracy: HomRank adopts a list-wise ranking that prioritizes the relative ordering of homogeneous candidates, rather than predicting absolute RMSD scores, enabling it to effectively distinguish fine-grained structural differences between high-fidelity conformations from low-fidelity ones. (3) Robustness to outliers: HomRank inherently reduces sensitivity to outliers, as the evaluation process emphasizes rankings rather than absolute scores.

Table 2 summarizes the comparisons of RMSD regression and HomRank.

**Table 2:** RMSD Regression vs. HomRank.

| Design | RMSD regression | HomRank |
| --- | --- | --- |
| Target | Scoring heterogeneous RNAs | Ranking homogeneous RNAs |
| Loss function | MSE loss | ListNet ranking loss |
| Input data in mini-batch | Heterogeneous RNAs & RMSD | Homogeneous RNAs & RMSD |
| Model output | RMSD (exact value) | Ranking order (relative value) |

### Architecture and Model Configurations

The architecture of HomRank is based on ARES [29], with key modifications including the design of a homogeneous RNA sampler to ensure homogeneous candidates within a mini-batch and the integration of the ListNet ranking technique to predict their ranking scores. The neural network is a sequential model comprising an atomic embedding layer to initialize atom representation, three blocks to capture structural equivariance and invariance, and three fully-connected layers to fuse representation and generate the ranking scores. Each block includes a self-interaction layer, an equivariant convolution layer, a pointwise normalization layer, followed by a self-interaction layer, and a non-linearity layer.

HomRank and HomRank-PT are trained on four Nvidia A100 GPUs, each equipped with 80GB of memory. The training leverages Distributed Data Parallel (DDP) to ensure synchronous gradient updates across GPUs. The batch size for each GPU is 32, with each mini-batch comprising 32 randomly sampled mini-lists *π*, each containing 10 candidates. The optimization employs the Adam optimizer with a learning rate 5*e*-3. The temperature coefficient *T* for the softmax transformation is fixed at 0.1. The threshold *δ* for ΔRMSD is set to 0.1Å. Consistent with the settings in ARES [29], the model is trained for a single epoch. For HomRank-PT, the model is optimized using the MSE loss during the pre-training stage, with batch size and learning rate aligning with [29]. In the training stage, the parameter configurations remain consistent with HomRank.

### Evaluation Metrics

#### (i) Minimum RMSD

Measures the minimum RMSD among the top-*k* evaluations:

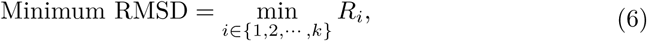

where {*R*_1_*, R*_2_, · · · *, R_k_*} represent the ground-truth RMSD corresponding to the top-*k* best-scoring candidates.

#### (ii) Mean RMSD

Measures the arithmetic mean RMSD among the top-*k* evaluations:

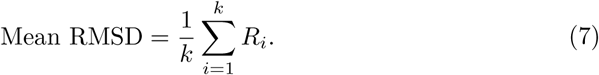

#### (iii) Success Rate (SRate)

The success rate SRate*_i_* for the *i*-th RNA, is a binary indicator defined as follows,

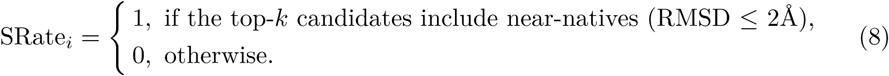

##### Average Success Rate 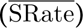

The average success rate is the arithmetic mean of SRate*_i_* across *m* RNAs, i.e., 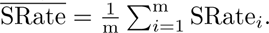

#### (iv) Retrieval Rate (RRate)

The retrieval rate RRate*_i_* for the proportion of near-native conformations retrieved within the top-*k* evaluations:

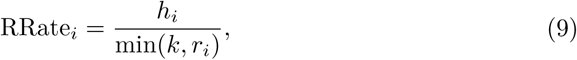

where *h_i_*denotes the number of near-native candidates retrieved within the top-*k* best-scoring candidates, *r_i_*is the total number of near-natives, and min(*k, r_i_*) accounts for cases where *r_i_* fewer than *k*.

##### Average Retrieval Rate 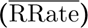

The average retrieval rate is the arithmetic mean of RRate*_i_* across *m* RNAs, i.e., 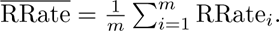

#### (v) Spearman’s rank correlation coefficient (*ρ*)

The Spearman’s rank correlation coefficient, *ρ_i_*, for the *i*-th RNA is a nonparametric measure of rank-based correlation, measuring the alignment between the ranking of candidates sorted by predicted score and the real ranking sorted by ground-truth RMSD. Let *n_i_* be the total number of candidates for the *i*-th RNA, and let *d_i,j_* be the difference in ranks for the *j*-th candidate. The coefficient *ρ* is calculated by,

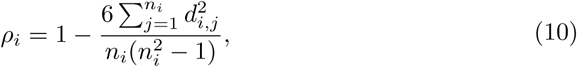

where a higher *ρ_i_*(upper bound is +1) indicates stronger alignment between the predicted ranking and the ground-truth ranking, while a lower value (lower bound is -1) implies a strong inverse relationship, and 0 denotes no correlation.

##### Average Spearman’s rank correlation coefficient (*ρ̄*)

The average Spearman correlation is the arithmetic mean of *ρ_i_* across *m* RNAs, i.e., 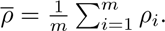

## Data Availability

The data supporting the findings of this study are available from the corresponding authors upon reasonable request. All input data are freely available from public sources. RNA structures for training and testing were collected from PDB at https://www.wwpdb.org/ftp/pdb-ftp-sites.

## Code Availability

The code of HomRank is available at https://github.com/liliangnudt/HomRank. Data were analyzed using Numpy v.1.25.2 (https://github.com/numpy/numpy) and Matplotlib v.3.7.1 (https://github.com/matplotlib/matplotlib). Structures were visualized by Pymol v.2.6.0 (https://github.com/schrodinger/pymol-open-source).

## Acknowledgments

The authors thank Tin Vlasšić for the valuable discussions on this work. This work was supported by the A*STAR GAP project (No. I23D1AG079).

## Author Contributions

Y.P., R.H. and M.S. initiated the project. Y.Y. and Y.P. proposed the ranking model for RNA structure evaluation. L.L., I.M. and M.K. helped refine the design of the ranking model. M.K. and R.H. prepared and constructed the new RNA structure data. L.L., Y.Y., I.M., Y.P., R.H. and M.S. designed the experiments. L.L. implemented the model, conducted the experiments and wrote the paper. Y.Y. and Y.P. revised the paper. I.M. proofread and proposed numerous edits for the paper. All authors reviewed and approved the final paper.

## Supplementary Materials

### Dataset Clustering Map

Figure S1 plots the detailed clustering map. Note that it includes a large set of 418 RNAs, but due to the significant computation expense of molecular dynamics simulations, a subset of 190 RNAs is used in this work. Actually, we performed molecular dynamics simulations for all of them, but only approximately half of these simulations were fully completed. Despite the limited RNA data, the data processing is rigorous for evaluating model generalization, and extensive experiments validate that HomRank demonstrates promising scoring accuracy.

**Fig. S1:**
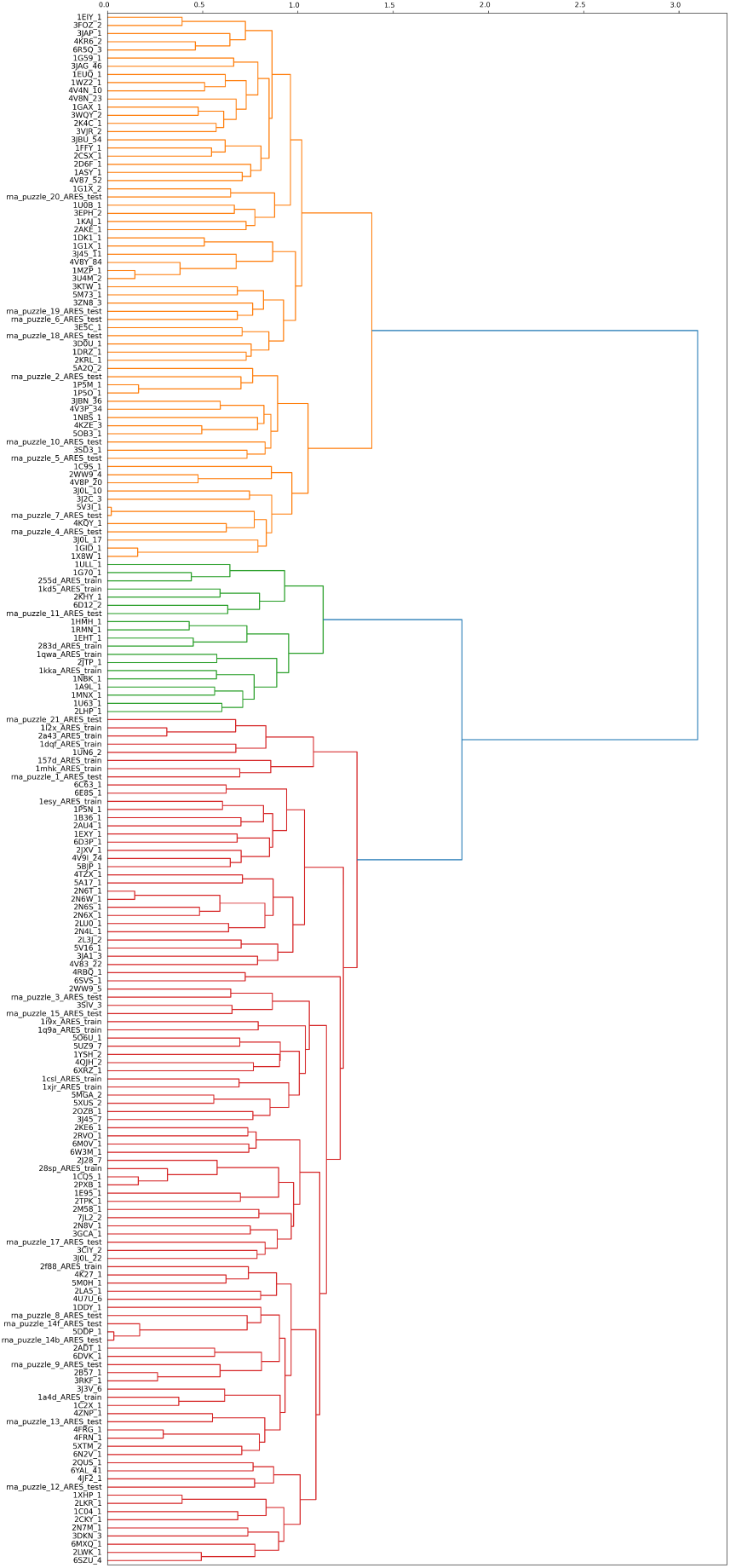
Clustering map of RNAs. The red (CLS-0), orange (CLS-1), green (CLS-2) clusters.

### Details of Testing Benchmarks

**Table S1:**
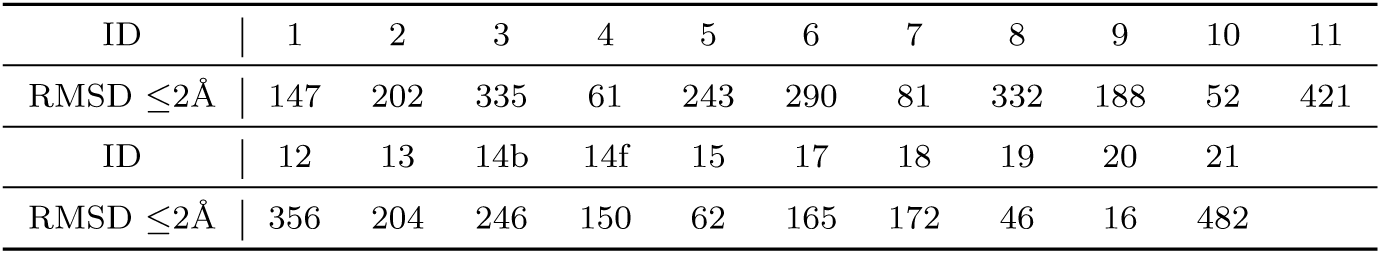
Number of near-native candidates in RNA Puzzles benchmark.

**Table S2:**
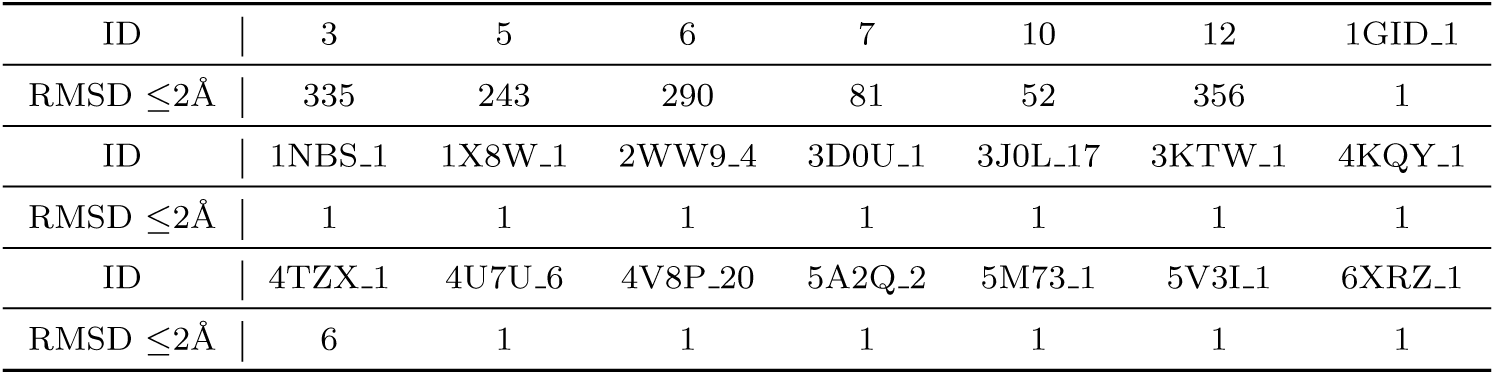
Number of near-native candidates in TestCLS-1 benchmark. The subscript is the entity ID.

**Table S3:** Number of near-native candidates in TestCLS-2 benchmark. The subscript is the entity ID.

|  |  |  |  |  |  |  |  |  |  |
| --- | --- | --- | --- | --- | --- | --- | --- | --- | --- |
| ID |  | 157D | 1CSL | 1DQF | 1ESY | 1KKA | 1MHK | 1QWA | 1XJR |
| RMSD $\leq 2\text{\AA}$ |  | 1 | 1 | 112 | 1 | 1 | 1 | 1 | 1 |
| ID |  | 255D | 283D | 1A9L.1 | 1B36.1 | 1EHT.1 | 1EXY.1 | 1G70.1 | 1HMH.1 |
| RMSD $\leq 2\text{\AA}$ |  | 40 | 1 | 1 | 1 | 1 | 1 | 1 | 1 |
| ID |  | 1NBK.1 | 1RMN.1 | 1U63.1 | 2AU4.1 | 2JTP.1 | 2LHP.1 | 6MXQ.1 |  |
| RMSD $\leq 2\text{\AA}$ |  | 1 | 1 | 1 | 1 | 1 | 1 | 1 |  |

### Ranking of Native Structures on the RNA Puzzles benchmark

Figure S2 depicts the ranking positions of native molecules using HomRank, HomRank-PT, and ARES+ in comparison with five baselines: ARES, Rosetta, cgRNASP, RNA3DCNN, and lociPARSE, on the RNA Puzzles benchmark. The diagonal blue line represents parity in rankings between the proposed model and the baseline, while points below this line indicate superior rankings by the proposed model. We note that HomRank and HomRank-PT consistently assign higher ranks to native conformations compared to the baselines, demonstrating the effectiveness of homogeneous RNA ranking in prioritizing near-native candidates. Additionally, ARES+ exhibits notable improvement over ARES.

**Fig. S2:**
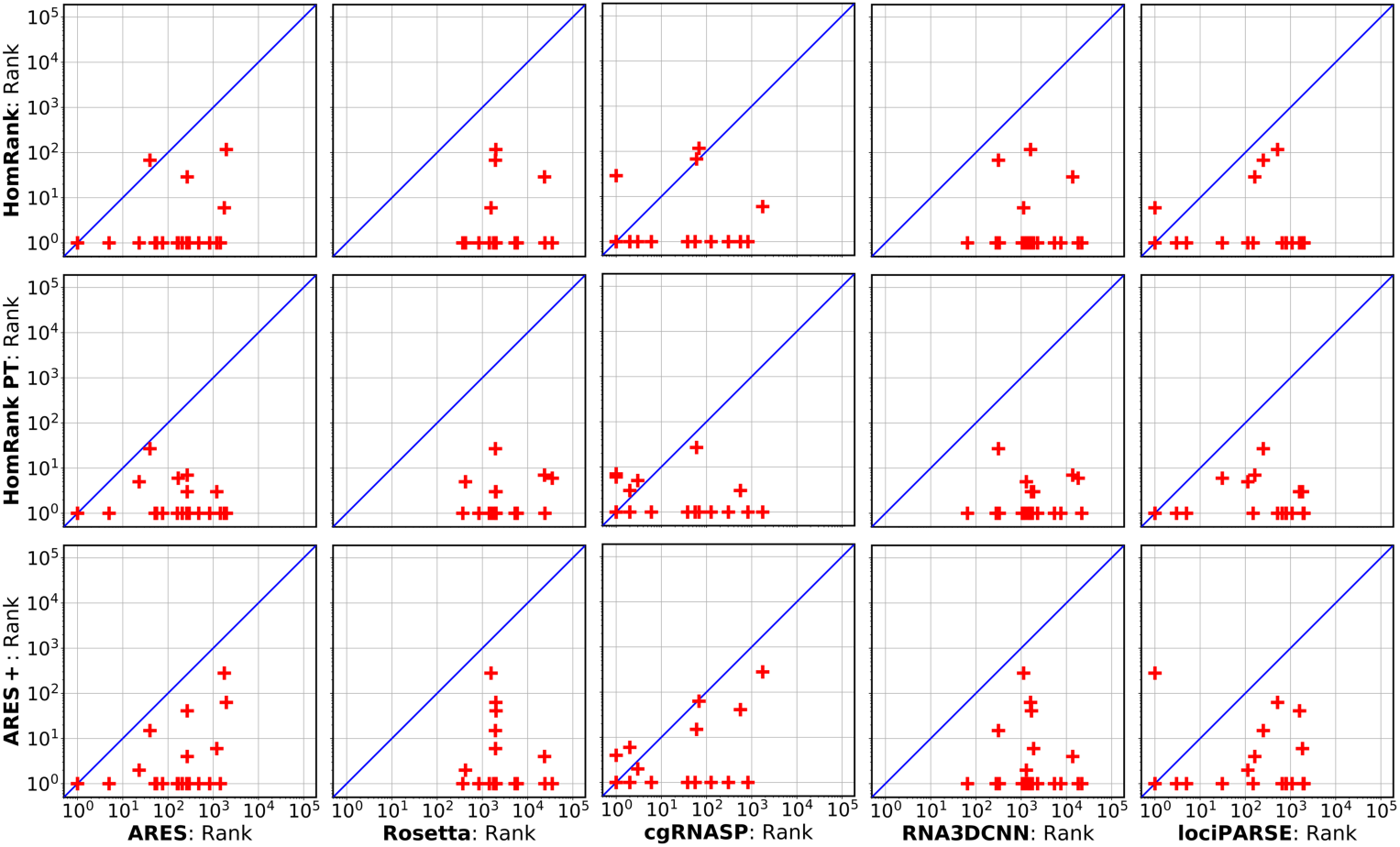
Ranking of native structure retrieval on the RNA Puzzles benchmark. This metric measures the ranking position of native RNA structures within the predicted score list.

### Experiments on the Six RNA Puzzles within TestCLS-1 Benchmark

Given that TestCLS-1 benchmark includes six RNA Puzzles that belong to different clusters from training data, we evaluate additional metrics on them.

Figure S3(a) measures the mean RMSD, a stricter metric than the minimum RMSD, as it reveals the average capacity of retrieving near-natives in the top 1, 10, 100 best-scoring candidates. The results show that HomRank and HomRank-PT significantly outperform both ARES and ARES+. Among the baselines, cgRNASP emerges as the strongest competitor, followed by lociPARSE. However, the widely used knowledge-based Rosetta scoring function and the pioneering deep learning-based RNA3DCNN, show poor performance on this metric.

**Fig. S3:**
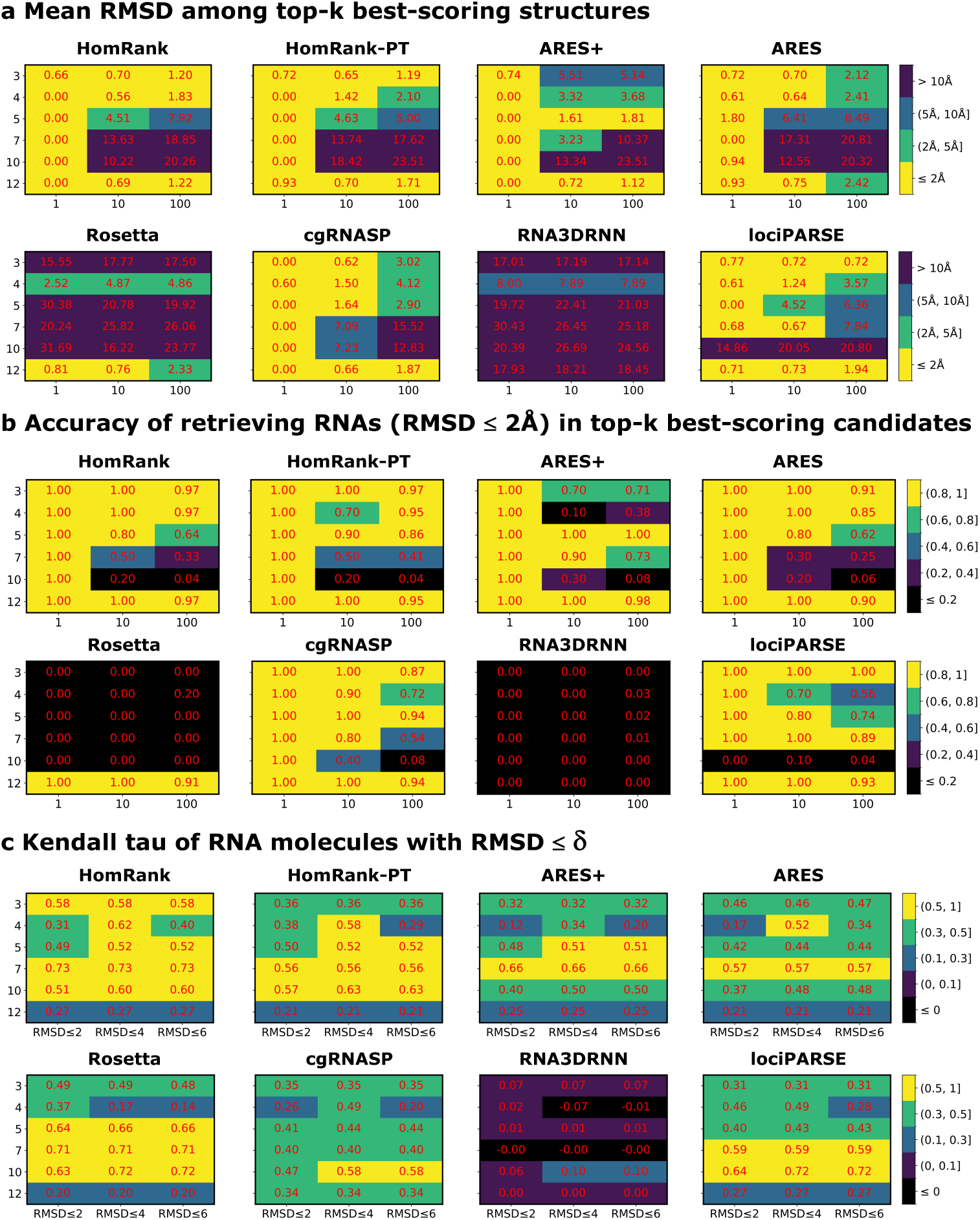
Evaluation on a subset of the TestCLS-1 dataset, containing six RNA Puzzles. **a.** Mean RMSD (red) of the top 1, 10, 100 best-scoring RNA molecules. **b.** Retrieval rate of retrieving near-natives within the top 1, 10, 100 evaluations. **c.** Kendall rank correlation coefficient *τ* for RNA molecules with RMSD ≤ 2Å, 4Å, 6Å.

It is worth noting that the number of near-native candidates varies among different RNAs, as illustrated in Table S1. Especially, RNA Puzzles 5, 7, and 10 have only 61, 81, and 52 candidates with RMSD ≤ 2Å, respectively, whereas RNA Puzzles 3, 6, and 12 contain 335, 243, and 356 near-natives. This imbalance impacts the evaluation of mean RMSD metric for the top-100 best-scoring candidates, leading to relatively weaker performance for RNA Puzzles 5, 7, and 10, where fewer near-native structures are available. To deal with this, Figure S3(b) presents the retrieval rate metric, which does not rely on of the total number of near-native candidates. The results indicate that HomRank and HomRank-PT achieve higher retrieval rate than ARES and ARES+. Among the baselines, cgRNASP and lociPARSE perform comparably to HomRank and HomRank-PT. However, Rosetta and RNA3DCNN perform noticeably worse, particularly RNA3DCNN, which performs poorly in retrieving near-native conformations.

Figure S3(c) plots the Kendall rank correlation coefficient *τ* (Kendall tau) for RNAs with RMSD ≤ 2Å, 4Å, and 6Å, which evaluates the ordinal association between the ground-truth and predicted rankings. *τ* measures the rank correlation between the ground-truth RMSD list and the corresponding predicted score list, which is formulated as 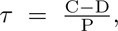 where C and D denote the number of concordant and discordant pairs, respectively, and P is the total number of pairs. *τ* = 1 means perfect agreement (all pairs are concordant), *τ* = −1 means perfect disagreement (all pairs are discordant), *τ* = 0 means no correlation (equal numbers of concordant and discordant pairs). We observe that HomRank achieves the highest Kendall tau metric in most cases, HomRank-PT exceeds ARES+ by large margins, while ARES+ performs comparably to ARES. These results demonstrate the superiority of the ranking technique over RMSD regression in accurately capturing the relative ranking of homogeneous RNAs. Interestingly, Rosetta demonstrates better Kendall tau metric compared to other baselines. However, its performance on minimum/mean RMSD and retrieval rate for RNAs (RMSD ≤ 2Å) in the top-*k* best-scoring candidates is noticeably worse. This indicates that Rosetta, based on knowledge-based statistical potentials, can model relative rankings but fail to prioritize high-fidelity candidates at the top of the ranking.

### Experiments on TestCLS-2 Benchmark

This section evaluates the performance on the TestCLS-2 benchmark. Figure S4 plots the top-*k* average success rate and average retrieval rate of retrieving near-native molecules. For the top-1 evaluation, all models exhibit relatively weak performance, with HomRank achieving the highest metrics, though still below 20%. As more candidates are considered, the metrics improve for all models, particularly for HomRank-PT, which demonstrates notable gains beyond the top-100 evaluations. Overall, HomRank, HomRank-PT, and ARES+ are the top three models for this benchmark, significantly outperforming ARES. Among the baselines, cgRNASP proves to be the most competitive, followed by RNA3DCNN and lociPARSE, while Rosetta shows the worst performance.

**Fig. S4:**
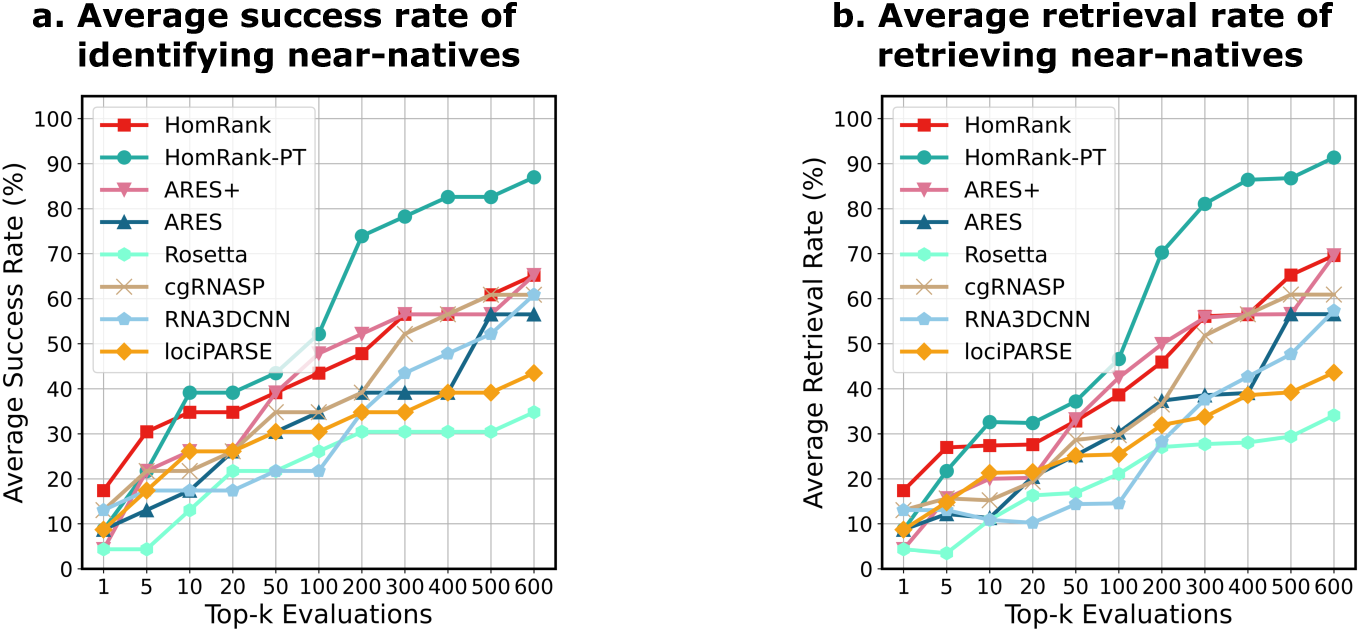
Comparisons on the TestCLS-2 benchmark. **a.** Average success rate across all RNAs in identifying native conformations within the top-*k* evaluations. **b.** Average retrieval rate across all RNAs for retrieving native conformations within the top-*k* evaluations.

### Case study

Figure S5 showcases the best-scoring candidates identified by each method. For the first case, 5A2Q 2, HomRank, HomRank-PT, and ARES+ successfully identify the native molecule, while the baselines mistakenly select decoy structures with significant RMSD deviations ranging from 15.70Å to 23.21Å. In the second case, 1GID 1, HomRank is the only model to accurately pinpoint the native conformation, showing the most competitive performance. Although HomRank-PT fails to discern the native one, it selects a higher-fidelity candidate (RMSD=5.44Å) than ARES+ (RMSD=6.52Å) and other baseline methods, whose RMSD spanning 5.45Å to 7.50Å.

**Fig. S5:**
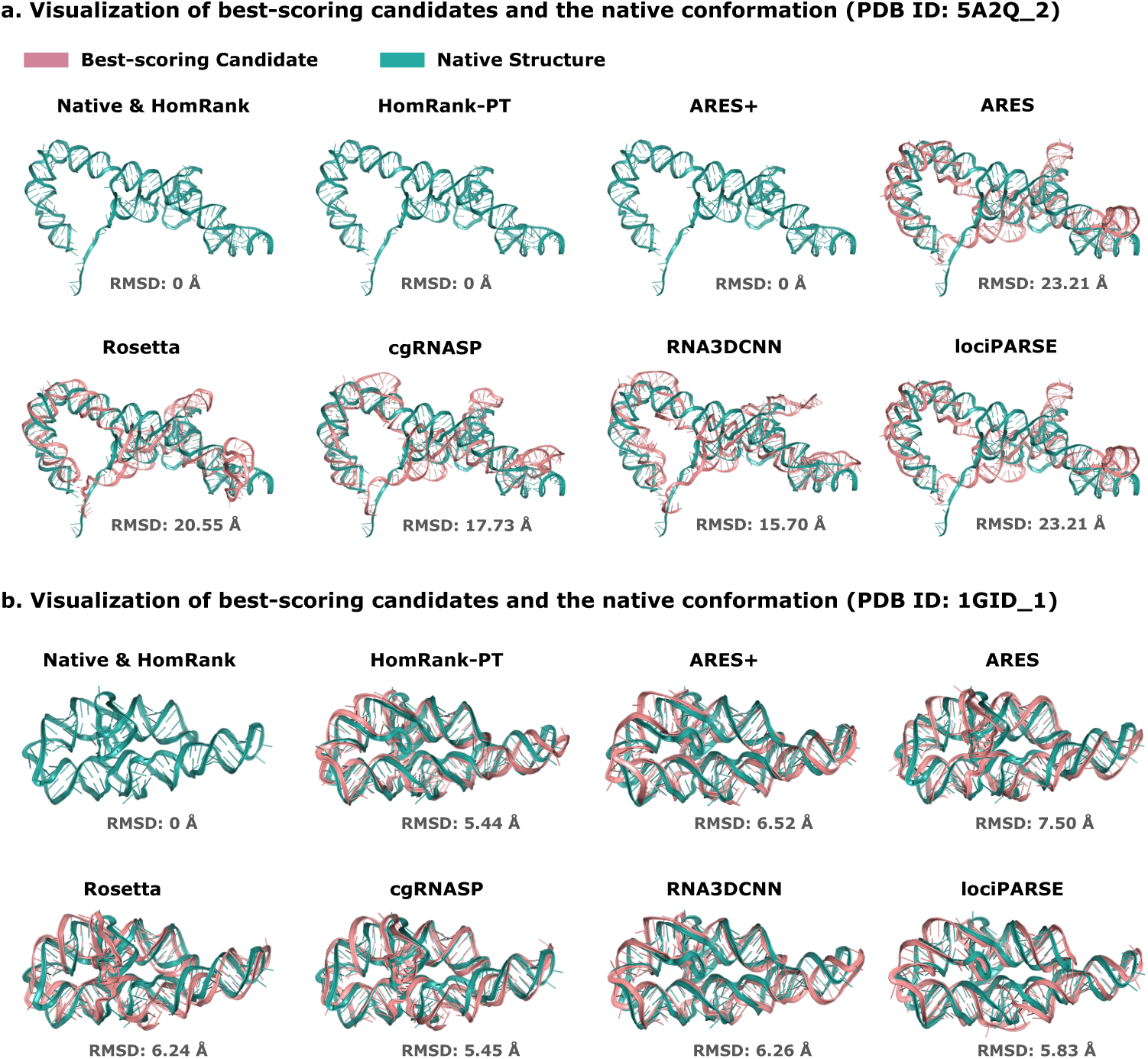
Case study. **a.** Example 1 (PDB ID: 5A2Q 2) demonstrates the effectiveness of HomRank, HomRank-PT, and ARES+ in accurately pinpointing the native molecule (RMSD=0Å), unlike the compared baselines that select decoy structures (RMSD: 15.70-23.21Å). **b.** Example 2 (PDB ID: 1GID 1) highlights HomRank as the only model capable of accurately identifying the native conformation (RMSD=0Å), whereas other models select decoy candidates (RMSD: 5.44-7.50Å).

## Footnotes

1 The 15 RNA Puzzles within CLS-0 are excluded from the training data, as these are commonly recognized as standard benchmarks [22, 29, 34]

